# The variation of promoter strength in different gene contexts

**DOI:** 10.1101/2020.07.21.214239

**Authors:** Tam T. Tran, Trevor C. Charles

## Abstract

**Background:** Promoter engineering has been employed as a strategy to enhance and optimize the production of bio-products. There have been many effortless studies searching the best promoter for biological application. However, whether promoter strengths stay unchanged in different gene contexts remains unknown.

**Results:** Six consecutive promoters at different strength levels were used to construct six different versions of plasmid backbone pTH1227, followed by inserted genes encoding two polymer-producing enzymes. Some of promoter strengths in the presence of inserted sequences did not correspond to the reported strengths in a previous study. When removing the inserted sequences, the strengths of these promoters returned to their reported strengths. These changes were further confirmed to occur at transcriptional levels. Polymer production using our newly constructed plasmids showed polymer accumulation levels relatively corresponding to the promoter strengths reported in our study.

**Conclusion:** Our study revealed the essence of re-assessing promoter strength in a specific gene context. Different gene contexts could result in the variation of promoter strengths, hence this might lead to different outcomes in downstream applications.

## Background

Promoter engineering has been considered as one of the critical strategies in the metabolic engineering field. A number of studies have been carried out to predict and characterize the promoter strength using *in silico* and/or wet-lab experiments [1–4]. Strong promoters are normally desired to maximize the end products, which could be proteins, amino acids, polymers or biofuel, and reduce toxicity during the growth phase. However, promoter strength and optimal yield of end products are not always correlated proportionally; therefore, promoters that are fine-tunable and tightly controlled are more beneficial and versatile.

The broad-host range expression vector pTH1227, a pFUS derivative vector, contains an RK2 origin, inducible pTac-*lacI*^*q*^ promoter and a reporter *gusA* gene [5–9]. This vector was used to study the role of the *minCDE* genes in both *S. meliloti* and *E. coli* [5]. The related vector pMP220 has been used in *Azorhizobium caulinodans* [6]. It was also used for insertional mutagenesis, transcriptional signal localization and gene regulation studies in root nodule bacteria such as *S. meliloti*, *Bradyrhizobium* sp. and *R. leguminosarum* bv. *Viciae* [8].

A library of engineered promoters of different strengths has been constructed through either mutagenesis of constitutive promoters or chemical synthesis of promoter variants [10–12]. By changing the length and the sequence of the spacer between the −35 and −10 regions, a broad range of promoters which differ in strength have been studied in *Lactococcus lactis.* Randomizing the spacers resulted in a remarkable change in activities up to 400-fold [11]. It was also emphasized that the context in which the consensus sequences are embedded significantly influences the promoter strength. In a subsequent study, this strategy has been employed to obtain the expression of the *pyrG* gene encoding CTP synthase at different levels and its effect on the growth rate and nucleotide pool size [10]. Another library of nearly 200 promoters was also obtained using error-prone PCR to examine gene expression in *E. coli* [12]. It was suggested that the optimization of gene expression could also depend on the genetic background of the strain.

To study gene expression or the effect of a specific promoter on gene expression, a reporter gene located downstream of the promoter is normally used. There are various reporter genes which have been employed to detect gene expression used in different host backgrounds such as *gfp* gene encoding green florescence protein or *gusA* gene from *E. coli* encoding β-glucuronidase [1, 2, 12–14]. The β-glucuronidase (Gus) product can be easily detected by observation of colour change on media containing chromogenic substrates such as X-GlcA (5-bromo-4-chloro-3-indolyl β-D-glucuronide, cyclohexyl-ammonium salt), other 3-indoxyl derivatives, and naphthol-β-D-glucuronide. In addition, it also can be quantitatively measured by spectrophotometry (p-nitrophenyl β-D-glucuronide and phenolphthalein-β-D-glucuronide), fluorimetry (4-methylumbelliferyl-β-D-glucuronide and 5-dodecanoyl-aminofluorescein-di-β-D-glucuronide) or chemiluminescence (1,2-dioxetane-β-D-glucuronide). Gus has been widely used to study gene expression and other applications because the enzyme is highly stable, resistant and easily detected. The *gfp* gene was used to examine gene expression in *E. coli* of nearly 200 promoters created by error-prone PCR [12]. These promoter strengths were also assessed by testing the effect of the expression of the *ppc* gene encoding phosphoenol pyruvate (PEP) carboxylase on growth yield and encoding deoxy-xylulose-P synthase on lycopene production.

In our study, we employed a broad range of constitutive promoters of different strengths and re-assessed their strengths in a specific context. Subsequently we investigated their effect on polymer production.

## Methods

### Strains and media used

All bacteria and plasmids were listed in Table 1. *S. meliloti* and *E. coli* strains were cultivated in Tryptone Yeast extract (TY) and Luria-Bertani (LB) media, repectively. Streptomycin (200 μg/ml) and tetracycline (10 μg/ml) were added if necessary.

**Table 1.** List of strains and plasmids were used in this study.

| Strains or plasmids | Genotype | Reference |
| --- | --- | --- |
| *E. coli* |  |  |
| DH5α | *supE44 lacU169(w80lacZDM15) hsdR17 recA1 endA1 gyrA96 thi-1 relA1* | Strain collection |
| *S. meliloti* |  |  |
| Rm1021 | Spontaneous Sm isolate of *R. meliloti* SU47 | Strain collection |
| SmUW499 | Rm1021 $\Delta phbC$ *exoF*:: Tn5 | Strain collection |
| SmUW255 | SmUW499 harboring pTH1227 | This study |
| SmUW256 | SmUW499 harboring pTAM | This study |
| Plasmids |  |  |
| pRK600 | pRK2013 npt::Tn9 Cm<sup>r</sup> Nm-Km<sup>s</sup> | [24] |
| pTH1227 | pFUS1 carrying *lacI*<sup>q</sup>-P<sub>tac</sub> DNA from pMal-c2x (AB32193/AB32194), Tc<sup>r</sup> | [5] |
| pTAM | pTH1227 *pct532 phaC1400* (codon-optimized) | [25] |
| pTAM2 | pTAM derivative, P<sub>0</sub> promoter- a substitute for *lacI*<sup>q</sup>-P<sub>tac</sub> | This study |
| pTAM3 | pTAM derivative, P<sub>1</sub> promoter- a substitute for *lacI*<sup>q</sup>-P<sub>tac</sub> | This study |
| pTAM4 | pTAM derivative, P<sub>2</sub> promoter- a substitute for *lacI*<sup>q</sup>-P<sub>tac</sub> | This study |
| pTAM5 | pTAM derivative, P<sub>3</sub> promoter- a substitute for *lacI*<sup>q</sup>-P<sub>tac</sub> | This study |
| pTAM6 | pTAM derivative, P<sub>4</sub> promoter- a substitute for *lacI*<sup>q</sup>-P<sub>tac</sub> | This study |
| pTAM7 | pTAM derivative, P<sub>5</sub> promoter- a substitute for *lacI*<sup>q</sup>-P<sub>tac</sub> | This study |
| pTAM2nopctphaC | pTAM2 derivative without *pct532 phaC1400* | This study |
| pTAM7nopctphaC | pTAM7 derivative without *pct532 phaC1400* | This study |

### GusA activity assay

This assay was carried out following the procedure as previously described [15]. Cells were fully grown in TY media, and then the culture was added into the assay buffer at the ratio of 1:4, incubating at room temperature until the mixture turned into a yellow color, at which point sodium carbonate was added to terminate the reaction. Reaction time and the absorbance of the mixture at 420 nm were recorded. Cell density of the culture was measured at 600nm for normalization.

### RNA isolation

RNases are present everywhere and very active, hence extra care must be taken to avoid sample degradation by using gloves and RNase-free materials. All solutions are prepared in RNase-free water or DEPC-treated water. Cells were sub-cultured in TY and grown to OD = 0.5 – 0.8. Stop solution which is 5% phenol in ethanol was added into the culture and mixed well to stop the cell growth. Next, culture was harvested by centrifuging at 5,000 rpm, 4°C for 10 min. Cells can be stored at −70°C after flash freezing in liquid nitrogen.

RNA extraction was performed following the hot phenol extraction protocol with some modification [16]. All the following steps should be carried out at 4°C, unless otherwise stated. The cell pellet from a 50 ml culture was thawed on ice and re-suspended in 960 μl of RNase-free water by vortexing. Next, 480 μl of hot phenol solution was added to an equal volume of cell re-suspension, vortexed vigorously and incubated in a water bath at 95°C for 1 min. The sample was spun down at 13000 rpm, 4°C for 10 min. Supernatant was added to 600 μl phenol/chloroform, and vortexed vigorously. Then, it was spun again for 5 mins, and aqueous phase was extracted twice with chloroform. To precipitate nucleic acid, the aqueous phase was added to 1/10 volume of sodium acetate and 2 volume of isopropyl alcohol, inverted to mix and incubated on ice for 30 min. Next, the sample was spun for 10min, washed with −20°C 70% ethanol and air-dried until the pellet turned translucent. Precipitate was then resuspended in 85 μl water and digested with 5 μl DNase I in 10 μl DNase I buffer at 37°C for 30 min to remove DNA from the sample. The DNA-free sample was confirmed by running on an agarose gel. If DNA is still present in the sample, water was added into the sample up to the volume of 400 μl, and nucleic acid precipitation was repeated as described above, followed by another round of DNA digestion. The RNA sample was further purified using Qiagen RNeasy mini kit, and finally resuspended in 100 μl water. The quantification of RNA was determined using the Nanodrop, and the ratio of A_260_/A_280_ should be around 2.0. The quality of RNA was checked by imaging the agarose-formaldehyde gel to observe rRNA bands.

### Dot blot

Samples with equal amount of RNA were applied onto a positively charged membrane using the Bio-Dot Microfiltration Apparatus (Bio-Rad). The membrane was placed on the gasket and wetted with 2x SSC. Saran wrap was used to cover unused wells. RNA samples were blotted to each well, followed by washing with 200 μl TE. Vacuum was used to help liquid pass through the membrane. Membrane fixation was performed using the UV-crosslinker instrument, using C-L program. Prehybridization was performed in DIG Easy Hyb buffer (Roche) at 40°C for 30-45 min, followed by hybridization overnight at the same temperature. Washing and detection steps were carried out as outlined in the manufacturer’s protocol for the DIG High Prime DNA Labeling and Detection Starter Kit II (Roche).

### Polyhydroxybutyrate (PHB) analysis

Polymer production was evaluated by gas chromatography following a protocol which has been described by others (Braunegg et al., 1978; Jung et al., 2009). Cell pellets were collected from flask culture after a 3-day incubation by centrifuging at 4,000 g for 20 min, then washed twice with distilled water, and finally dried at 100°C overnight. The dried cell weight (DCW) was recorded before methanolysis in 2 ml chloroform and 1 ml PHB solution containing 8 g benzoic acid l^−1^ as an internal standard and either 30% sulfuric acid (for PHB) in methanol. The reaction was carried out at 96°C for 6 h, cooled, and then 1 ml of water was added, the mixture was vortexed, and the solution was allowed to separate into two phases. 1 μl of the chloroform phase was taken for analysis by GC as previously described (Jung et al., 2009). The samples were injected into Agilent 6890 series GC system with a DB Wax column (30 m × 0.53 mm, film thickness 1 μM, J & W Scientifics), which is located in the Department of Chemical Engineering. The oven program was set as following: initial temperature was set at 80°C for 5 min, then ramped to 230°C at 7.5°C/min, and continued to ramp to 260°C at a faster rate 10°C/min followed by maintaining that temperature for 5 min.

## Results

### Construction of plasmids with promoters of different strengths

A range of promoters of different strengths (6 out of 12 native promoters) based on a previous study [2] was chosen for further investigation. These promoter sequences (Table 2) were synthesized in the form of extended primers that were annealed to create double-strand DNAs. The backbone plasmid, pTH1227a, is a derivative of pTH1227 that had the XbaI site removed by being digested, blunted and self-ligated as shown in Fig. 1. Promoter P_0_ was designed to contain XbaI site next to HindIII site and cloned into pTH1227a at the HindIII and XhoI sites, creating pTH1227b. Next, the synthesized fragment which contains two engineered genes were inserted into the pTH1227b at the XhoI and PstI sites. The rest of the promoter set containing BglII for post-cloning confirmation was cloned into XbaI and XhoI, replacing promoter P_0_. As a result, we have constructed 6 different versions of the pTAM plasmid carrying different promoters of varying strengths.GusA activities obtained by expressing *gusA* gene under a set of constitutive promoters of different strengths in the presence of synthesized genes

**Table 2.**
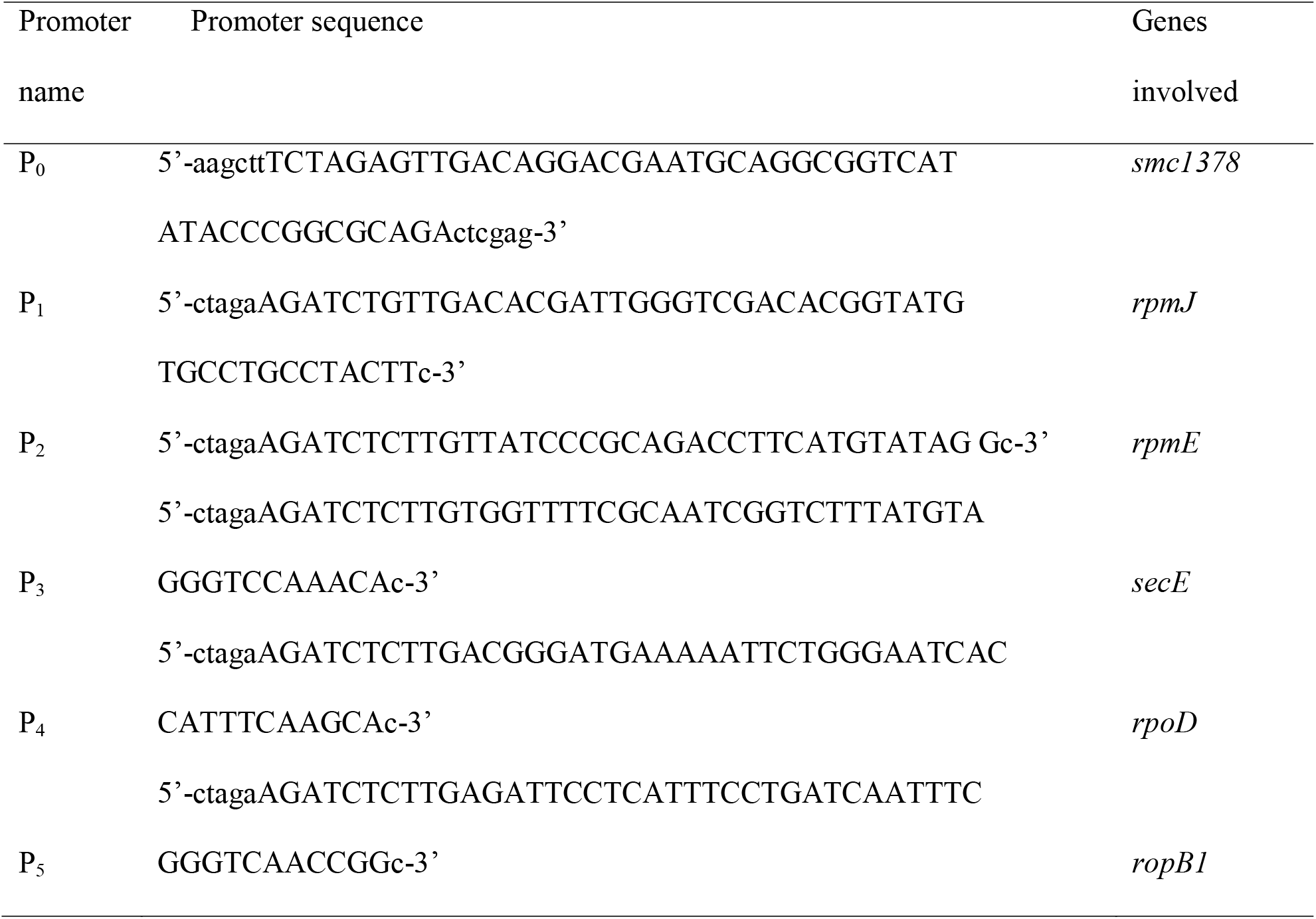
List of native promoters, their sequences and original genes which were expressed under these promoters in *S. meliloti*

**Fig. 1.**
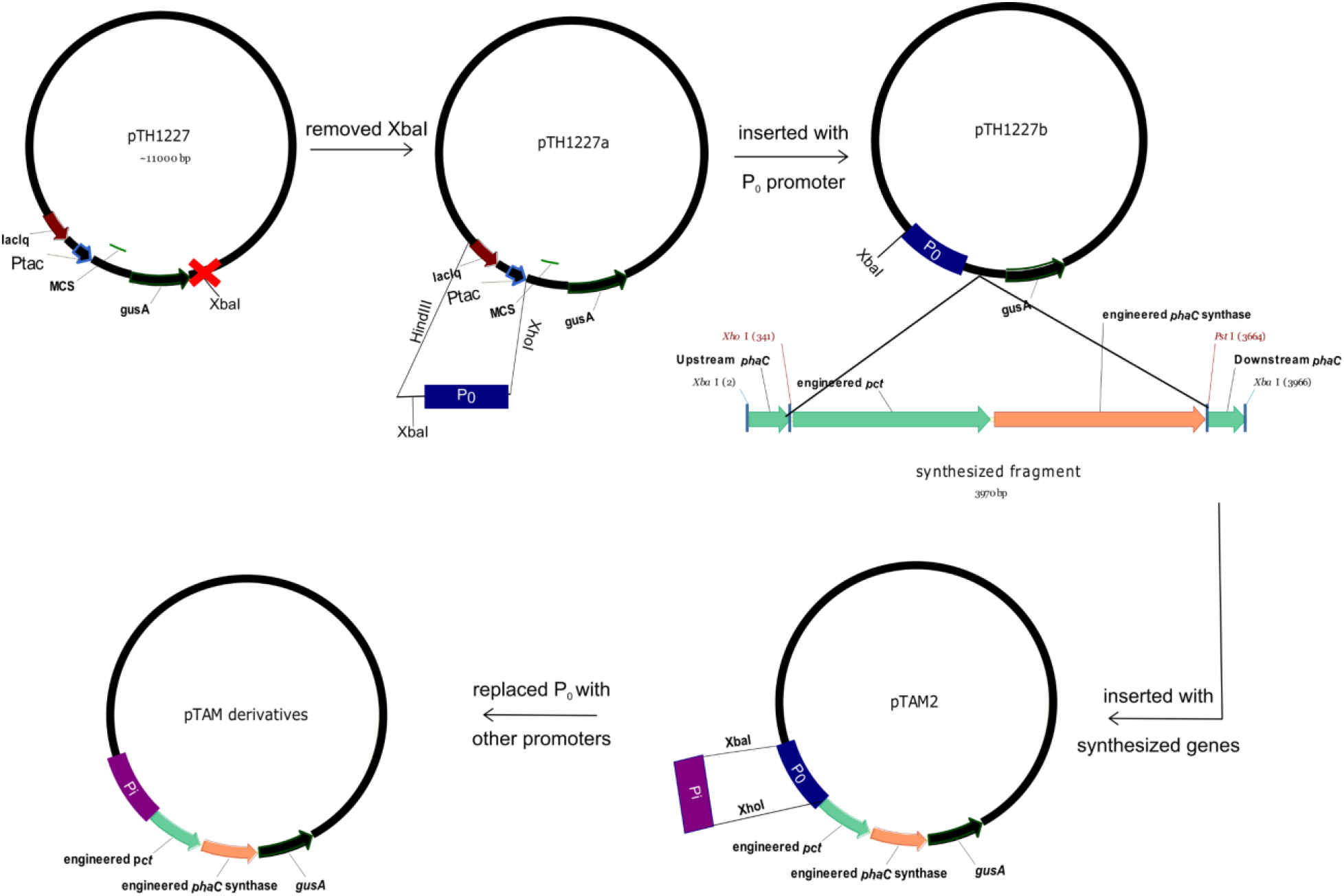
Construction of plasmids with promoters of different strengths. In the last step, P0 was replaced with Pi which is one of five promoters (P1, P2, P3, P4 and P5).

GusA activity assays were performed to examine the level of gene expression according to each specific promoter that was used. As shown in Fig. 2, a wide range of GusA activity was obtained indicating that different levels of gene expression were successfully achieved. Also, in some cases the level of gene expression was even lower than the background level of gene expression obtained by using the inducible promoter.

**Fig. 2.**
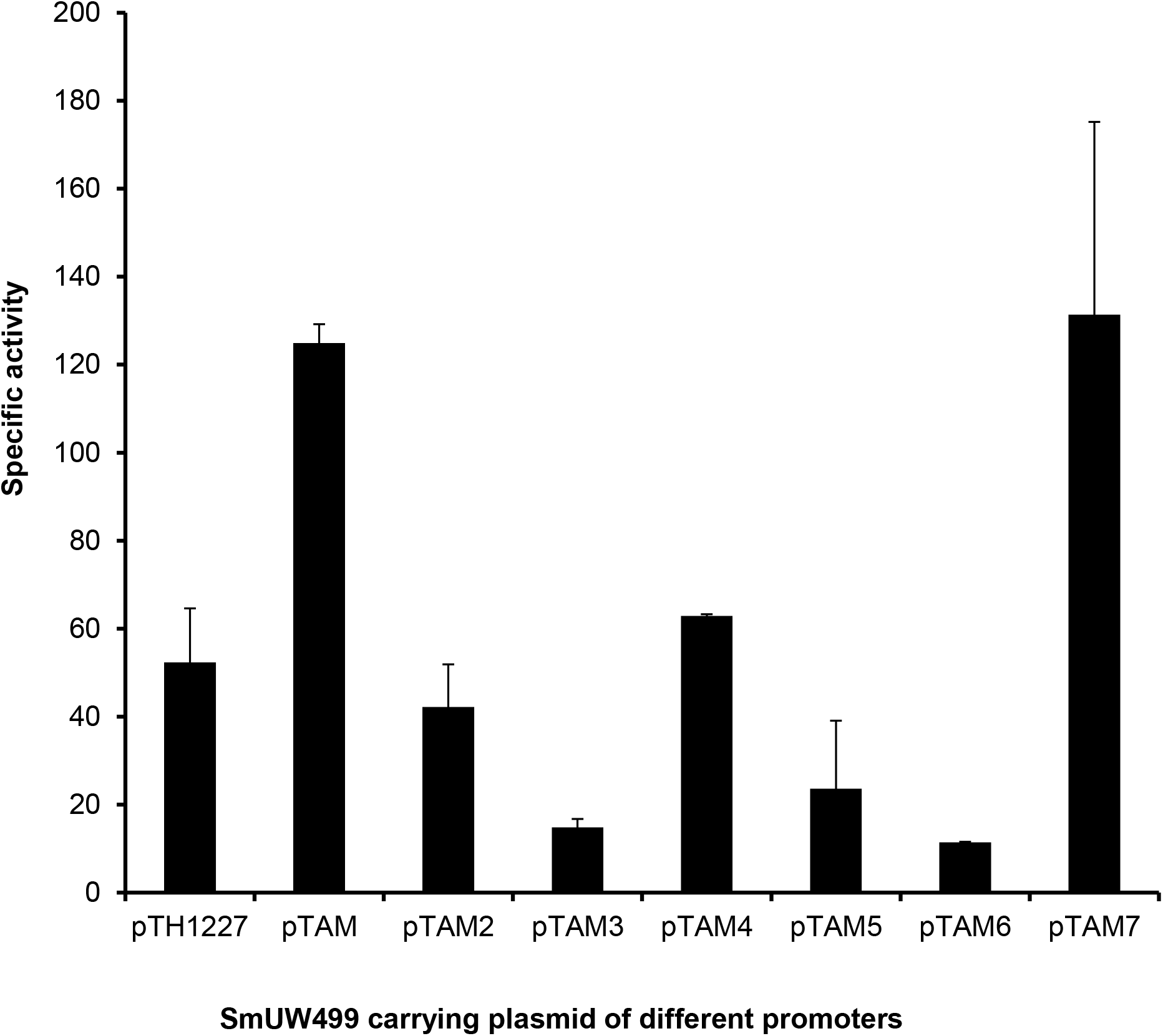
GusA specific activity of *S.meliloti phaC* mutant SmUW499 harboring either the parental plasmid (pTH1227) or different pTAM derivatives (pTAM, pTAM2 -> pTAM7). pTH1227: empty plasmid, inducible pTac promoter; pTAM: inducible pTac promoter + insert genes; pTAM2: *smc1378* gene promoter + insert genes; pTAM3: *rpmJ* gene promoter + insert genes; pTAM4: *rpmE* gene promoter + insert genes; pTAM5: *secE* gene promoter + insert genes; pTAM6: *rpoD* gene promoter + insert genes; pTAM7: *ropB1* gene promoter + insert genes.

### GusA activities obtained by expressing *gusA* gene under a set of constitutive promoters of different strengths in the absence of synthesized genes

To further investigate the effect of the synthesized genes on the promoter strength, we chose three plasmids, pTAM2, pTAM6 and pTAM7, which carried the *smc1378, rpoD* and *ropB1* promoters, respectively. Since the expression from the *smc1378* and *ropB1* promoters was in strong disagreement with the previous study of MacLellan et al. on expression of these reporter genes while *rpoD* gene maintained the weakest activity in both our study and the previous study, we thought that it would be interesting to examine the expression of the reporter gene under these promoters in the absence of the intervening synthesized genes. Therefore, the fragment containing the synthesized genes was removed by XbaI and PstI digestion followed by self-ligation, named pTAM2no*pctPhaC*, pTAM6no*pctPhaC* and pTAM7no*pctPhaC*. Surprisingly, removal of synthesized genes recovered the original promoter activities which were reported in by MacLellan et al. for strains which have genes expressed under the *smc1378* and *ropB1* promoters (Fig. 3). The *rpoD* promoter still maintained the low activity which was observed with and without the inserted genes.

**Fig. 3.**
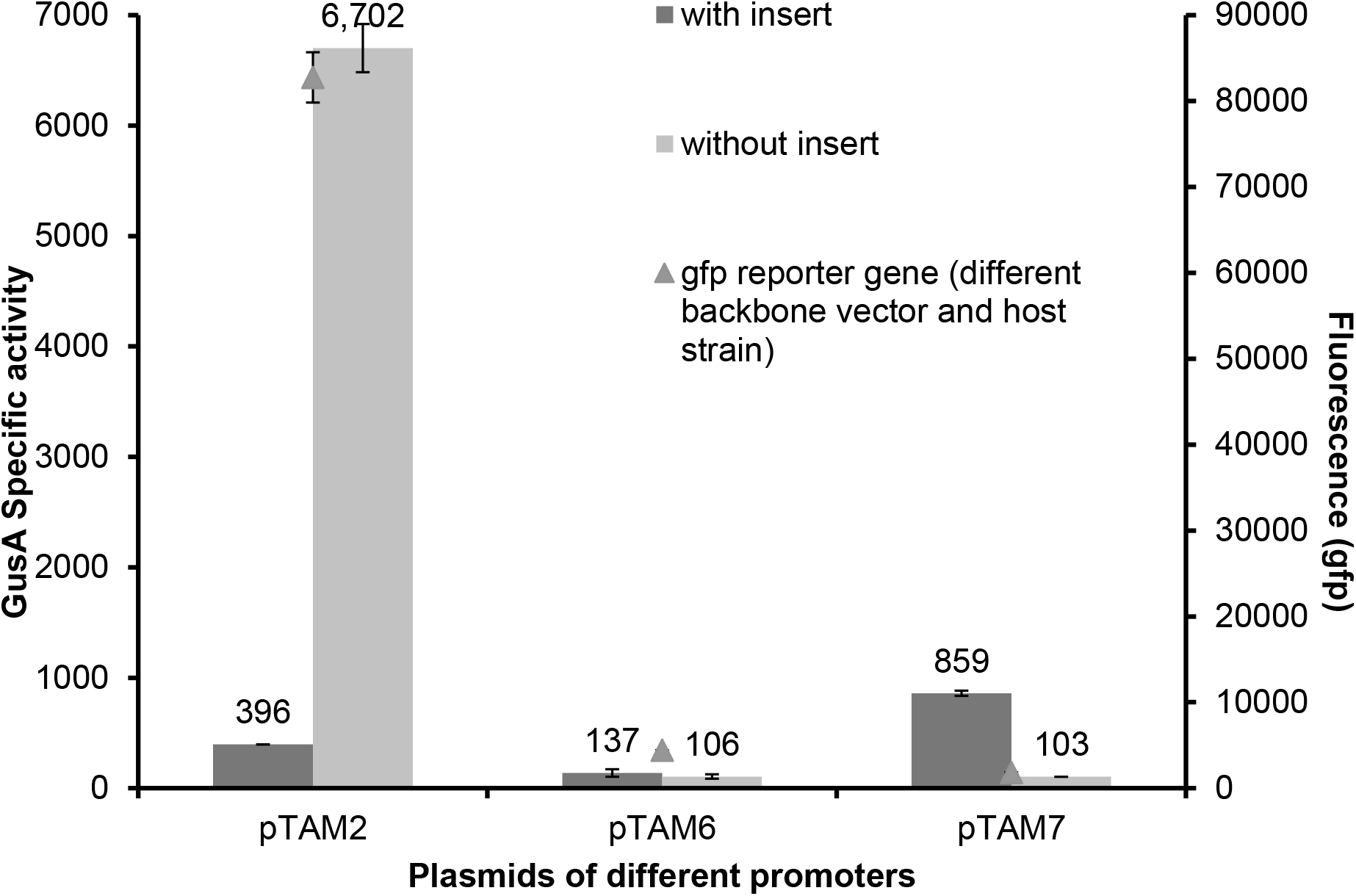
Comparison of activity of *gusA* gene (left axis) and *gfp* gene (right axis). Dark bars: *S. meliloti* strains SmUW499 carrying one of three plasmids pTAM2, pTAM6 and pTAM7 which are composed of *smc1378, rpoD*, *ropB1* gene promoters, respectively, and synthesized genes were measured the activity of *gusA* gene. Light bars: *S. meliloti* strains SmUW499 carrying one of three derivative plasmids pTAM2, pTAM6 and pTAM7 which are composed only *smc1378, rpoD*, *ropB1* gene promoters, respectively, were measured the activity of *gusA* gene. Triangle dots: the activity of *gfp* gene which were expressed under the corresponding promoter have been reported in a previous study (MacLellan et al., 2006)

### Analysis of transcript abundance of *gusA* and *pct* genes using dot blot

From the above results, the presence of synthesized genes that changed genetic context has shown the effect on the activity level of some promoters which were indirectly assessed by measuring *gusA* activity of the reporter gene. As shown in Supplementary 1 and Fig. 4, the abundance of *gusA* transcripts corresponded with the GusA activity, suggesting that the presence of synthesized genes caused the decrease and increase of *gusA* transcripts for pTAM2 and pTAM7 plasmids, respectively. The abundance of *pct* transcripts was also investigated and compared to that of *gusA* transcripts on the plasmids carrying the synthesized genes (Supplementary 2). As our expectation, the abundance of *pct* transcripts is similar to that of *gusA* transcripts since they are transcriptionally fused.

**Fig. 4.**
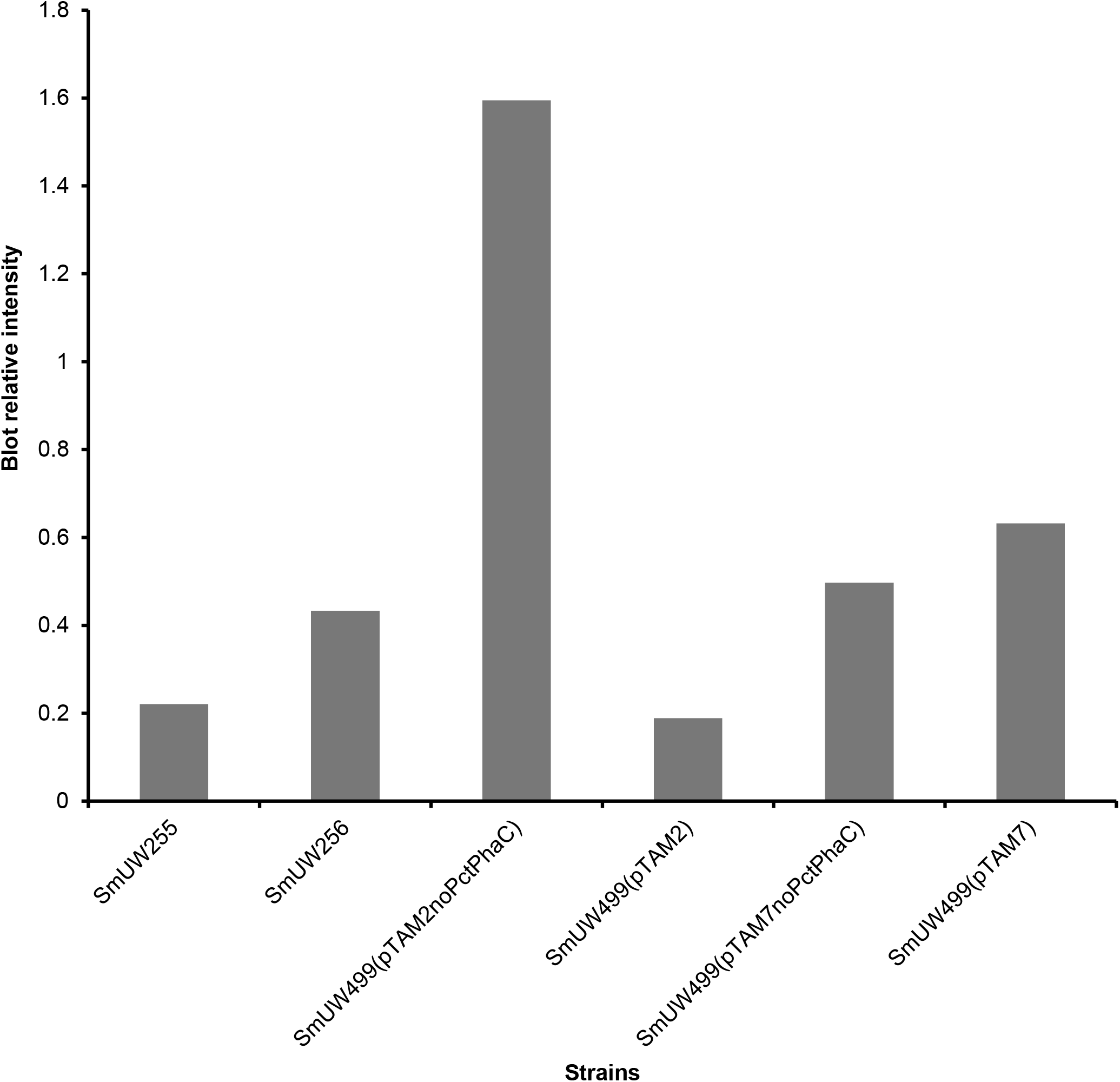
Graph represents the relative intensity on dot blot results as shown in Supplementary 1

### Polymer production in strains that have synthesized genes expressed under these constitutive promoters

To evaluate the different expression profiles on polymer production, these strains were cultivated in define media supplemented with mannitol as a substrate and grown under polymer accumulating conditions. As expected, strains carrying pTAM6 and pTAM7, which had the weakest and strongest promoter strengths produced the least and most amount of PHB, respectively (Fig. 5). The strains carrying pTAM7 which showed promoter strength similar to that of the strain carrying the *tac* promoter under induction conditions also accumulated a similar amount of PHB. Other strains showing a range of intermediate promoter strengths produced similarly low amount of polymer.

**Fig. 5.**
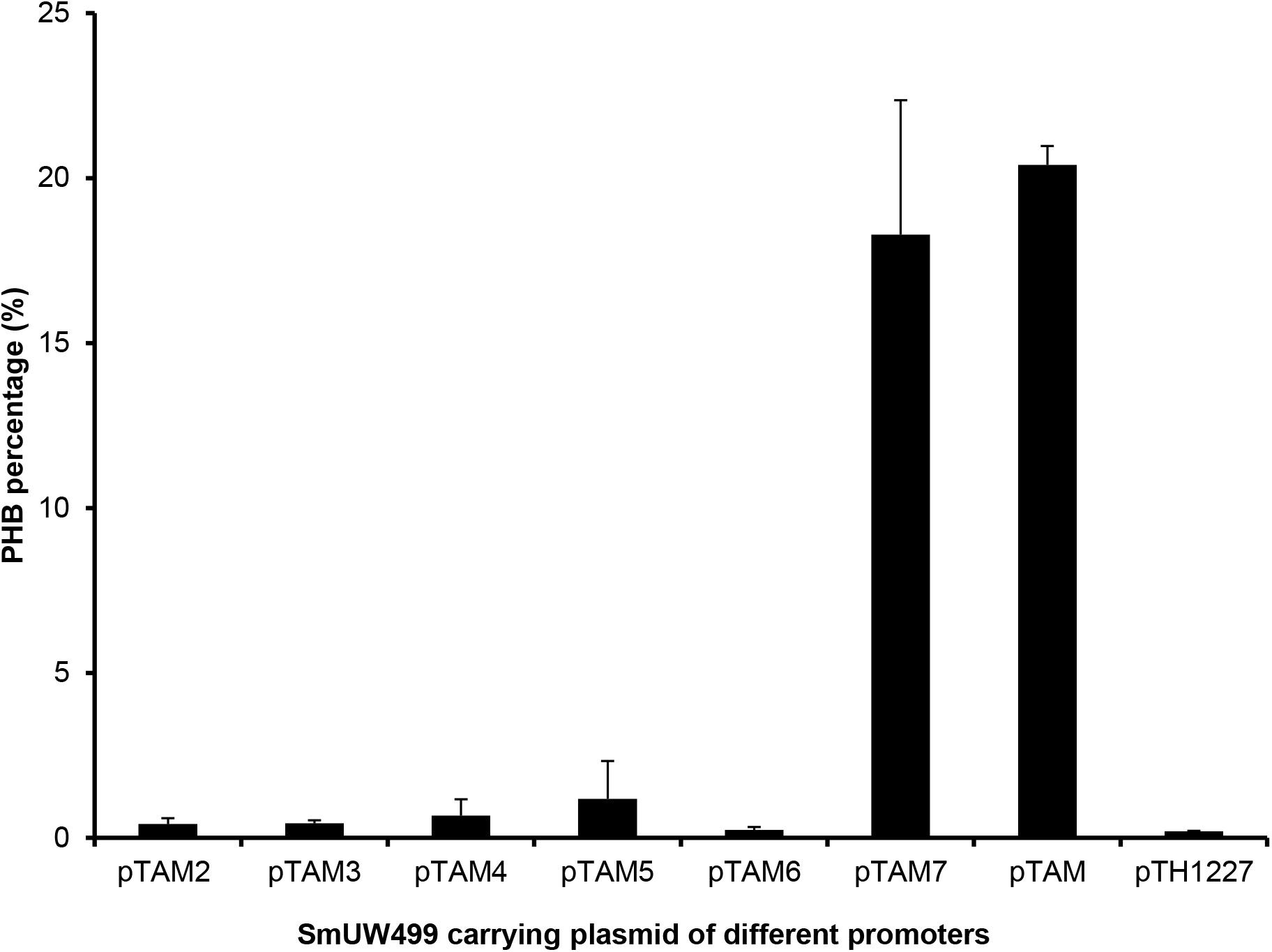
PHB production in *S.meliloti phaC* mutant strains SmUW499 that have synthesized genes are expressed under different native promoters (pTAM2 -> pTAM7) and inducible promoter (pTAM); the negative control *S.meliloti* SmUW499 strains carrying the empty plasmid pTH1227

## Discussion

The transcriptional levels of some promoters in our study were completely contradictory to previously reported values. For example, we found that the pTAM7 plasmid carrying the *ropB1* promoter exhibited the highest levels of expression, while MacLellan et al [2] had previously demonstrated that this same promoter had the lowest level of expression among the same set of promoters. Another promoter that strongly disagreed with MacLellan et al in terms of gene expression level was the *Smc1378* gene promoter in the pTAM2 plasmid. In the previous study, expression was the highest of all of the promoters, but we found that the expression was relatively low compared to other promoters in the same set. This finding raised concerns about potential context effects on gene expression. It could be due to plasmid backbone, inserted gene sequence, the reporter gene employed or the cultivation condition. In a previous study, the relative promoter activity has been proved to remain unchanged across different cultivation conditions [17]. The activities of roughly 900 *S. cerevisiae* and 1800 *E. coli* promoters are taken into consideration, and their gene expressions changed among different cultivation conditions by a constant factor. In other words, they mostly behaved alike and maintained their relative activities levels across different cultivation conditions. Therefore, the cultivation condition could be ruled out as a possible cause. Among the remaining potential causes, we postulated that inserted gene sequence most probably caused the irrelative change in gene expression of some promoters. Therefore, we removed this inserted sequence from the plasmid construct and compared the GusA activity of the construct with and without these inserted genes.

As our postulation, removing the insert bought back the promoter strength reported in that previous study. This suggested that the downstream sequence probably influenced the transcription process. Other work has shown that the downstream region can have a strong influence on the efficiency of the escape process of RNA polymerase because release of the σ subunit only occurs after the polymerase has transcribed 8-11 nucleotides [18, 19]. This region was earlier reported by Bujard and co-workers [20]. They found that the down-stream sequence could change the promoter strength *in vivo* more than 10-fold. The reason is that RNA polymerase covers a larger region than 35 bp (up to 70 bp); hence, its activity also depends on the flanking regions which could be up-stream or down-stream regions. It was known that this region was involved in promoter escape; however, the mechanism of this process was not understood. By further study of the anti initial transcription sequence (ITS) which was discovered earlier to have an effect on promoter strength, it was found that the function of the anti ITS did not depend on either the stability of RNA:DNA bond or the interaction with core RNA polymerase. It appeared that the function of anti ITS was related to the σ subunit. As a result, the interactions between them could influence the promoter escape. In other studies, it was also observed that characterized promoters often showed variable activities depending on the genetic locus or gene transcribed [11, 12, 21–23]. A library of variable-strength, constitutive promoters was designed and constructed in bacteria [4]. The length of these promoters is 160 bp including extended sequences at both the 5’- and 3’-ends, so-called insulation sequences. Their promoter strengths were shown to maintain constant relative levels and be independent of genetic contexts. From the above evidence, we reason to suspect that the initial sequence of roughly 20 bp in the synthesized gene fragment may have had some effect on promoter strengths. Since the promoter activities were indirectly assessed through the expression of *gusA* reporter gene, there is no clear evidence to prove that these observations happened at the transcriptional level, not translational level. Therefore, we continued to investigate the abundance of RNA transcripts of *pct* and *gusA* genes in the pTAM2 and pTAM7 plasmids to verify these observations occurring at the transcriptional level.

The abundance of *gusA* and *pct* transcripts were directly proportional to GusA activity. These results support the hypothesis that the downstream sequence influenced on the transcription process which led to the change in the abundance of transcripts and enzyme activity as a consequence. However, which sequence of the downstream region and how far is it from TSS are still not identified. As mentioned earlier, the first 20 amino acids following the TSS might play an important role in either facilitating or impeding the transcriptional process. Therefore, it would be interesting to further investigate these sequences in the future. There have been a number of studies focusing on the effect of the −10 and −35 regions on promoter strength. However, only the upstream region before the transcription start site (TSS) has been seriously taken into consideration. In our study, it has been shown experimentally that this adverse effect exists. In the presence of engineered genes, relative promoter strength is different from what was reported. Removal of the insert fragment restored their relative strengths consistently across different reporter genes. We also demonstrated that the change occurred at the transcriptional level of both insert fragment and reporter gene.

## Conclusions

We selected a range of promoters of different strength that were well-characterized in the literature. However, our study did not agree with previous study, in which the relative promoter strengths have varied not correspondingly. We found that the inserted gene located close to promoters had a significant effect on promoter strengths. We postulated that this effect response may be due to effects on the occurrence during the releasing process of RNA polymerase from the σ subunit.

## List of abbreviations

PHB: Polyhydroxybutyrate

## Authors’ contributions

TT conceptualized the research idea and experimental analysis. TT designed and drafted the manuscript. TC revised the drafted manuscript and made necessary corrections. The authors read and approved the final manuscript.

## Acknowledgements

We thank Ricardo Nordeste, University of Waterloo, for the gift of strain SmUW499

