## Supplementary for "The variation of promoter strength in different gene contexts"

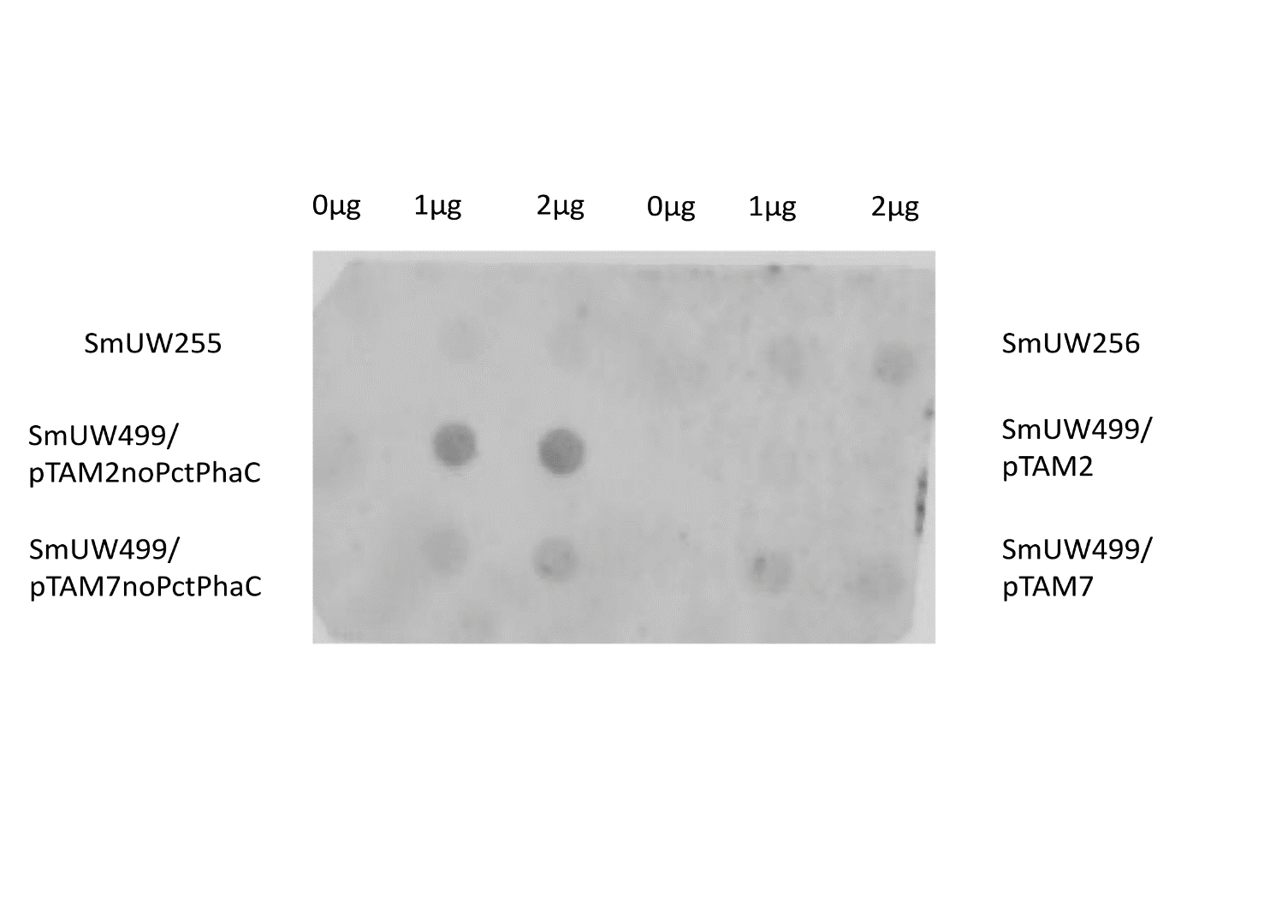

Supplementary 1 Dot blot image of *S. meliloti* SmUW499 carrying plasmids of different promoters using *gusA* probe. Three left columns represent RNA samples of different amounts (0, 1, 2μg) of SmUW255, SmUW499 (pTAM2no*pctphaC*), SmUW499 (pTAM7no*pctphaC*). The other right three columns represent RNA samples of different amounts (0, 1, 2μg) of SmUW256), SmUW499 (pTAM2), SmUW499 (pTAM7).

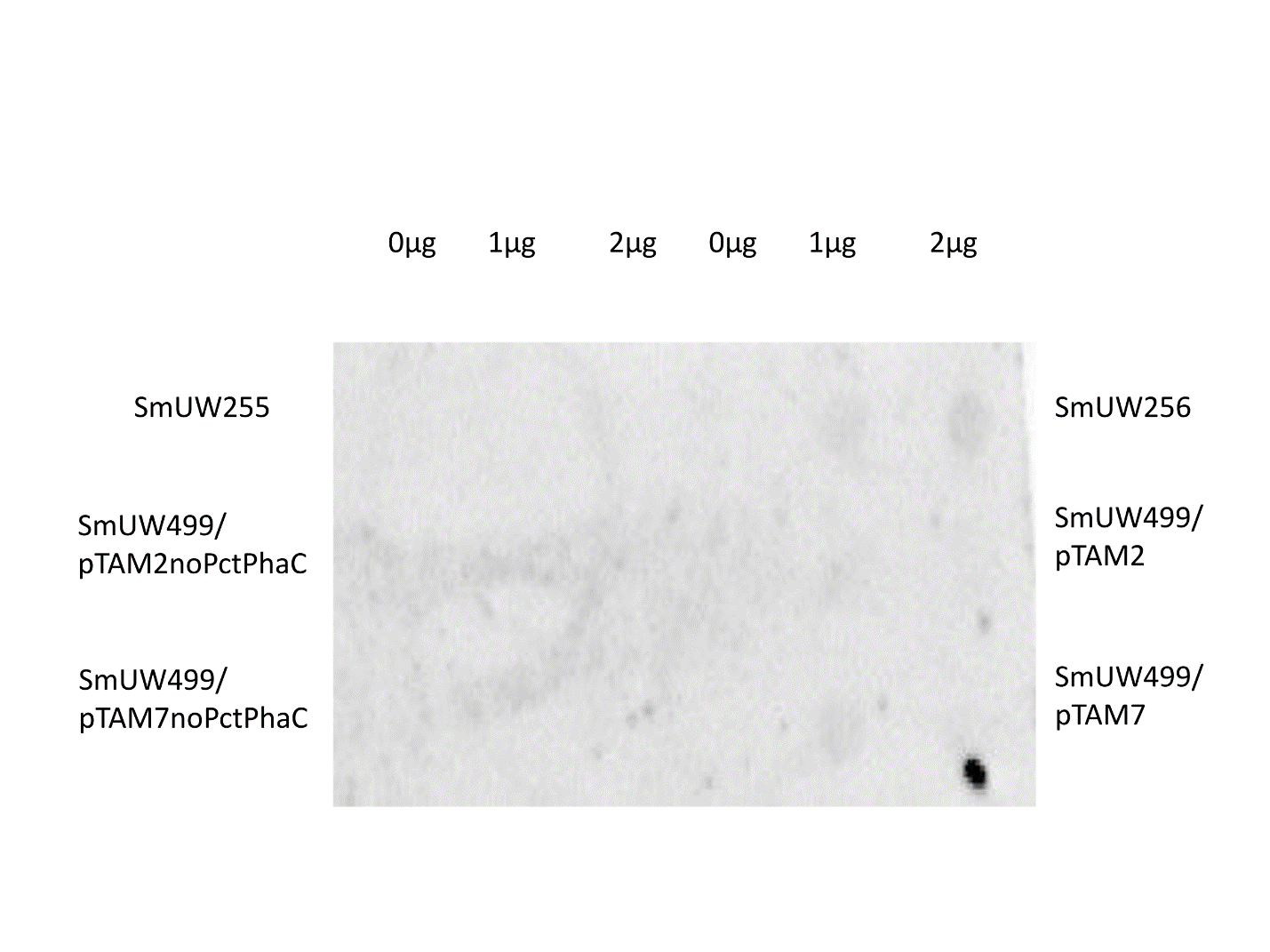

Supplementary 2 Dot blot image of *S. meliloti* SmUW499 carrying plasmids of different promoters using *pct* probe. Three left columns represent RNA samples of different amounts (0, 1, 2μg) of SmUW255, SmUW499 (pTAM2no*pctphaC*), SmUW499 (pTAM7no*pctphaC*). The other right three columns represent RNA samples of different amounts (0, 1, 2μg) of SmUW256), SmUW499 (pTAM2), SmUW499 (pTAM7).
